# Graph Contextualized Attention Network for Predicting Synthetic Lethality in Human Cancers

**DOI:** 10.1101/2021.01.27.428345

**Authors:** Yahui Long, Min Wu, Yong Liu, Jie Zheng, Chee Keong Kwoh, Jiawei Luo, Xiaoli Li

## Abstract

**Motivation:** Synthetic Lethality (SL) plays an increasingly critical role in the targeted anticancer therapeutics. In addition, identifying SL interactions can create opportunities to selectively kill cancer cells without harming normal cells. Given the high cost of wet-lab experiments, in silico prediction of SL interactions as an alternative can be a rapid and cost-effective way to guide the experimental screening of candidate SL pairs. Several matrix factorization-based methods have recently been proposed for human SL prediction. However, they are limited in capturing the dependencies of neighbors. In addition, it is also highly challenging to make accurate predictions for new genes without any known SL partners.

**Results:** In this work, we propose a novel *graph contextualized attention network* named GCATSL to learn gene representations for SL prediction. First, we leverage different data sources to construct multiple feature graphs for genes, which serve as the feature inputs for our GCATSL method. Second, for each feature graph, we design node-level attention mechanism to effectively capture the importance of local and global neighbors and learn local and global representations for the nodes, respectively. We further exploit multi-layer perceptron (MLP) to aggregate the original features with the local and global representations and then derive the feature-specific representations. Third, to derive the final representations, we design feature-level attention to integrate feature-specific representations by taking the importance of different feature graphs into account. Extensive experimental results on three datasets under different settings demonstrate that our GCATSL model outperforms 14 state-of-the-art methods consistently. In addition, case studies further validate the effectiveness of our proposed model in identifying novel SL pairs.

**Availability:** Python codes and dataset are available at:

## 1 Introduction

Human cancer is a variety of complex diseases that are commonly induced by the defects of multiple genes. Thus identifying genetic interactions is an essential step for anticancer drug development. Specifically, Synthetic Lethality (SL) is a type of genetic interaction, which occurs when the loss of either gene is viable while the loss of both is lethal. Recently, synthetic lethality has attracted increasing attention in cancer treatment since synthetic lethal interactions can be used to identify anticancer drug targets, and provide an opportunity to selectively kill cancer cells without harming the normal cells (Hartwell *et al.*, 1997; Iglehart and Silver, 2009). Therefore, cancer therapies based on the SL concept can result in fewer adverse effects compared with traditional chemotherapies (O’Neil *et al.*, 2017). More recently, high-throughput wet-lab screening techniques, such as small-molecule libraries (Chan *et al.*, 2011), RNA interference (RNAi) (Luo *et al.*, 2009) and CRISPR (Du *et al.*, 2017), have been developed to detect SL interactions. However, due to limitations such as high cost, off-target effects and unclear mechanisms of the web-lab screening, there is an urgent need to develop computational methods to complement the experimental screenings for SL interactions in human cancers.

In the past decade, some computational methods for predicting SL interactions have been developed. We can divide these methods into two main categories, namely *knowledge-based methods* and *supervised machine learning methods*. Knowledge-based methods are based on prior knowledge or hypotheses to predict SL interactions. For example, Jerby-Arnon *et al.* (2014) developed a data-driven model called DAISY for SL interaction identification by analysing gene expression and mutation data, based on the assumption that SL genes are often co-expressed but seldom co-mutated. Similarly, Sinha *et al.* (2017) predicted potential SL partners using mutation, copy number and gene expression data. Zhang *et al.* (2015) proposed to combine a data-driven model with knowledge of signaling pathways to identify SL interaction pairs by simulating the influence of gene knock-down to cell death. Apaolaza *et al.* (2017) attempted to use gene expression data to predict SL partners relying on the concept of minimal cut sets (MCSs), which refers to minimal sets of reactions whose removal would disable the functioning of a specific metabolic task. In addition, Jacunski *et al.* (2015) presented a network-based method to identify SL pairs by using the concept of connectivity homology, a type of biological connectivity patterns that persist across species. Srihari *et al.* (2015) introduced a novel in silico approach to identify SL interactions based on copy-number and gene expression data. However, knowledge-based methods depend strongly on prior knowledge and do not exploit the underlying patterns of known SL interactions to predict novel SL pairs.

With the rapid development of machine learning, supervised learning methods have been widely applied for various biological tasks, such as drug-target prediction (Liu *et al.*, 2016; Ezzat *et al.*, 2019; Zhang *et al.*, 2020), drug-microbe prediction (Long *et al.*, 2020), and gene-disease prediction (Natarajan and Dhillon, 2014). Recently, increasing supervised learning methods have been proposed for the SL prediction task. For example, Das *et al.* (2019) developed a random forest classifier-based prediction model named DiscoverSL to identity SL interactions in cancers using multi-omic cancer data (i.e., mutation, copy number alternation and gene expression data from TCGA). After that, Benstead-Hume *et al.* (2019) incorporated PPI and Gene Ontology (GO) data as feature sources and then predicted novel SL interactions also using the random forest classifier. Note that the above methods require manual extraction of different features for genes from diverse data sources. By modeling the SL data as a graph, graph embedding methods (e.g., matrix factorization) can automatically learn the gene features/embeddings in the SL graph for SL prediction. For example, Liu *et al.* (2019) first utilized logistic matrix factorization to learn representations of genes, and then used the learned representations to infer potential SL interactions. Liany *et al.* (2020) developed a novel prediction model for SL interactions based on collective matrix factorization (CMF) by simultaneously modeling multiple matrices describing the relation between the same entity. In addition, Huang *et al.* (2019) proposed a graph regularized self-representative matrix factorization-based method to identify SL partners. Unfortunately, all the aforementioned methods do not use the valuable topological structure information (i.e., neighbourhoods) of genes in the SL graph. To address this issue, Cai *et al.* (2020) presented a novel graph convolutional network (GCN)-based model named DDGCN to predict SL interactions. However, it has two limitations as follows. First, it predicts novel SL pairs based solely on the known SL data and does not utilize other data sources for feature extraction (e.g., it uses the SL matrix directly as gene features). Second, different neighbors contribute distinct importance to the centre node. Nevertheless, DDGCN considers the neighbors equally and is thus not able to preserve the importance of different neighbors.

In contrast, graph attention networks (GATs) (Veličković *et al.*, 2018) can effectively distinguish and preserve this kind of difference among neighbors, while capturing the graph structure and modelling the dependencies between neighboring nodes. GATs aim to aggregate the feature information of neighbors by assigning different weight values to different neighbors and has been successfully applied for various link prediction tasks in the fields of bioinformatics, such as microbe-disease prediction (Long *et al.*, 2020) and enhancer–promoter prediction (Hong *et al.*, 2020). Given that SL prediction is also a link prediction problem, we are thus motivated to generalize GAT for novel SL prediction. Nevertheless, there are still some challenges against employing GAT for SL prediction. First, while various data sources for human genes are now publicly available, it remains a challenge to effectively integrate them in a GAT framework for SL prediction. Second, many genes have no experimentally confirmed SL partners in current SL databases, e.g., SynLethDB (Guo *et al.*, 2016). We denote such genes without known SL partners as *new* genes. With no training data for *new* genes, it is very challenging for the GAT-based model to predict their SL partners.

To address the above challenges, we propose a novel Graph Contextual Attention Network called GCATSL for SL prediction. First, we exploit diverse biological data sources to construct multiple feature graphs for genes, which would serve as the feature inputs for GCATSL. Second, we design a dual attention mechanism, i.e., node-level attention and feature-level attention, to learn node (gene) representations from multiple feature graphs. Specifically, we first design a graph attention network with node-level attention to learn preliminary representations for nodes from each feature graph. To fully integrate the node representations from multiple feature graphs, we further implement feature-level attention to learn the importance of different feature graphs. Extensive experiments under different cross-validation settings (e.g., the setting to predict SL interactions for new genes) show that our model outperforms the state-of-the-art methods consistently. Case studies for top-predicted SL pairs further demonstrate the effectiveness of our GCATSL model. Overall, our main contributions are summarized as follows.

- We propose a novel GAT-based framework named GCATSL, which effectively incorporates various biological data sources for SL prediction. To the best of our knowledge, this is the first attempt to adapt GAT for SL prediction.
- We design a new dual-attention mechanism in our proposed GCATSL framework to effectively capture the different importance of neighbors and different feature graphs for node/gene representation learning.
- Comprehensive experiments are conducted on three datasets under different cross-validation settings (e.g., the setting for new genes). Experimental results demonstrate that our proposed GCATSL model outperforms 14 existing methods for identifying novel SL pairs.

## 2 Methods

In this section, we first describe the preliminaries and the problem formulation. Then we introduce our proposed GCATSL model in details.

### 2.1 Preliminary

Assuming that SL interactions are represented as a graph 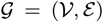 where 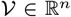 denotes the set of *n* nodes (i.e., genes), while *ε* denotes the set of edges (i.e., SL interactions). We denote *A* ∈ ℝ^*n×n*^ as the adjacency matrix of this graph, in which the entity *A*_*ij*_ is equal to 1 if gene *ν*_*i*_ is confirmed to have an SL interaction with gene *ν*_*j*_, otherwise 0. Moreover, we represent the set of observed/known SL interactions as Ω^+^ = {(*ν*_*i*_, *ν*_*j*_) ∈ Ω|*A*_*ij*_ = 1} with Ω denoting the set of all the gene pairs. Therefore, the unknown pairs can be represented as Ω^−^ = Ω\Ω^+^.

In this work, we derive the feature matrices/graphs for genes from different data sources (e.g., GO and PPI). For example, we calculate a semantic similarity matrix for genes based on their GO terms. We can thus consider this *n* × *n* similarity matrix as a feature graph. We denote each feature graph as ***H*** = {***h***_1_, ***h***_2_, …, ***h***_*n*_}, in which ***h***_*i*_ ∈ ℝ^*n*^ represents the feature vector of gene *ν*_*i*_ with the initial feature dimension *n*. To reduce dimension and extract more valuable features, we further implement Principal Component Analysis (PCA) on each feature graph/matrix to reduce the feature dimension from *n* to *d*_1_.

Our main task is to predict novel SL interactions based on the extracted gene features, as well as the graph for the observed SL interactions. Here we cast this important task as a link prediction problem in the SL graph. Specifically, with the observed SL interaction set Ω^+^ in the SL graph 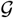 and the derived gene features *H*, we aim to learn a prediction function *f* : Ω → [0, 1] to infer the probability of an unknown pair in Ω^−^ to be a real SL interaction. Subsequently, according to the predicted scores, we prioritize all the pairs to determine the most possible candidate partners for a given gene.

### 2.2 Overview of GCATSL

In this paper, we propose a novel framework of Graph Contextualized Attention Network called GCATSL for SL prediction. As shown in Fig.1 (a), GCATSL mainly consists of three steps to learn gene representations and predict novel SL pairs. First, we learn feature-specific representations for each node with node-level attention from individual input feature graph respectively, as shown in Fig.1 (b). Second, we implement feature-level attention to aggregate node representations by assigning greater weights to more important feature graphs. Third, we reconstruct an SL interaction network based on the learned gene representations for SL prediction. Next, we introduce the three steps in details.

**Fig. 1.**
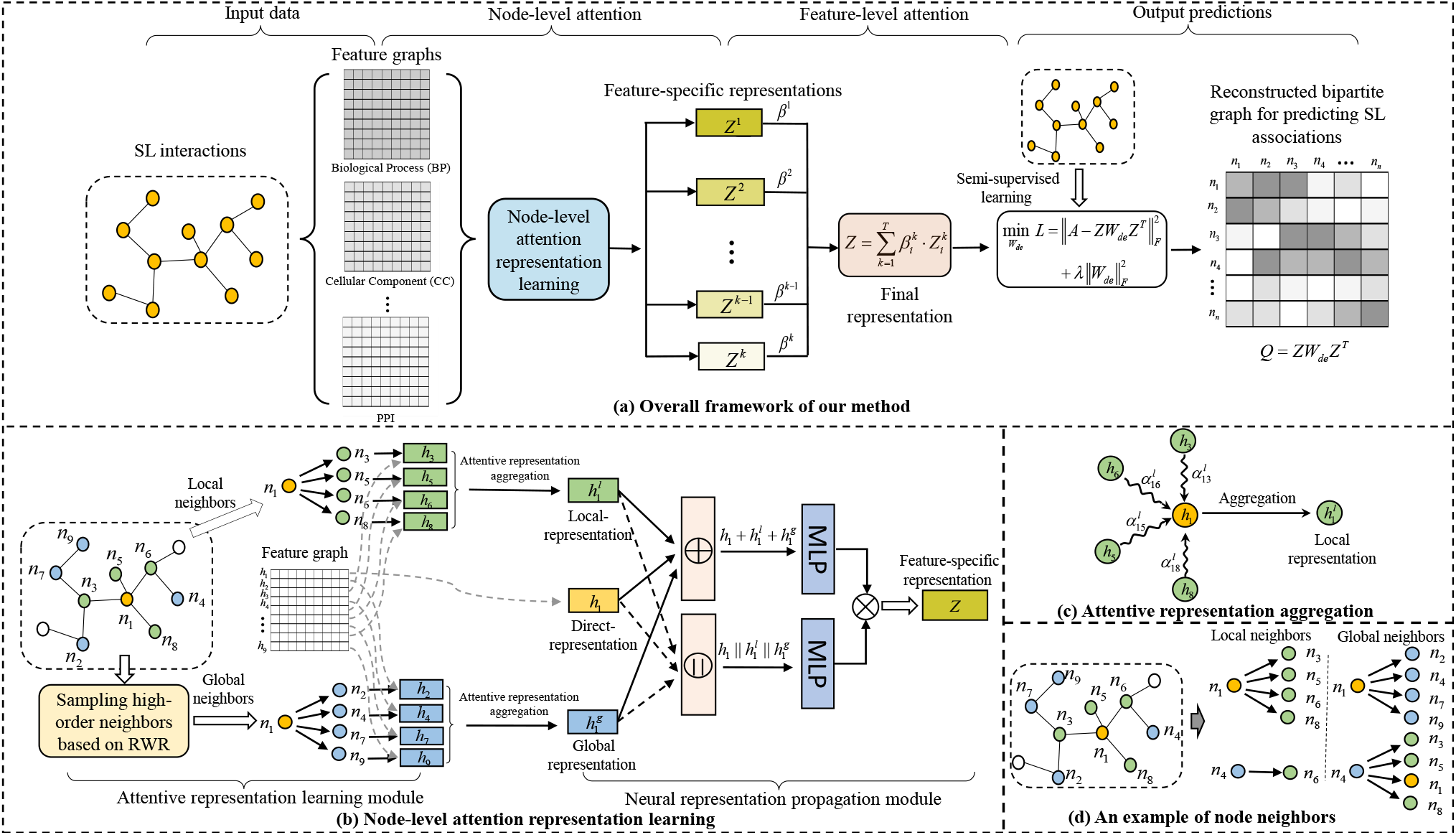
The overall architecture of GCATSL for gene representation learning and SL prediction. (a) Overall framework of our method. (b) Node-level attention representation learning. (c) Attentive representation aggregation. (d) An example of node neighbors.

### 2.3 Node-level attention for representation learning

In a specific feature graph, we notice that for each node, different neighbours play different roles and contribute different importance in learning node representation. To capture the importance, we introduce node-level attention to learn graph-specific representations for nodes. Given a feature graph ***H*** = {***h***_1_, ***h***_2_, …, ***h***_*n*_}, we first utilize the graph attention networks (GATs) (Veličković *et al.*, 2018) to learn the node representations from local neighbors and global neighbors in the SL graph 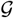. As shown in Fig.1 (b), we further design artificial neural networks (ANNs) to aggregate the node representations derived from different contexts. As the aggregated representation here is tied to the input feature graph ***H***, we thus denote it as feature-specific representation.

#### 2.3.1 Local representations

For a given node, we define the nodes that are directly connected to it in a graph as its local neighbors. For example, as shown in the left part of Fig.1(b), the local neighbors of *n*_1_ include nodes *n*_3_, *n*_5_, *n*_6_ and *n*_8_. Considering that different neighbors may yield biased importance, we design a node-level attention mechanism to learn the representations from local neighbors as shown in Fig.1 (c). Specifically, given a node *ν*_*i*_, we first learn the importance of its local neighbors by the following attention scores *e*_*ij*_.

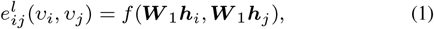

where 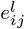 is the attention score that indicates the importance of the neighbor *v*_*j*_ to *v*_*i*_. In addition, *f* (·) denotes a single-layer feed-forward neural network and 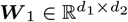 is a learnable weight matrix, which transforms input raw features with dimension *d*_1_ into high-level features of dimension *d*_2_ for genes.

To make the attention scores in Eq.1 comparable across different nodes, we further normalize them across 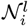, the set of all the local neighbors of node *v*_*i*_, and calculate the coefficient 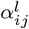 using the following softmax function:

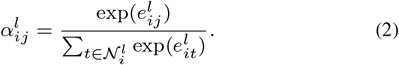

Subsequently, we derive the local representation 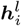 for node *ν*_*i*_ by aggregating the representations of its local neighbors according to their attention coefficients as follows.

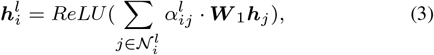

where *ReLU* (rectified linear unit) is the activation function.

Due to the instability of attention coefficient, individual node attention is likely to introduce noises. To reduce the noises, we further extend the node attention to multi-head attention by repeating the node attention for *K* times. We then concatenate the *K* learned representations into node *v*_*i*_’s local representation as follows:

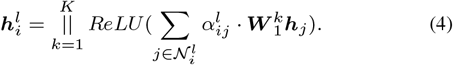

where || denotes the operation of vector concatenation.

#### 2.3.2 Global representations

Local representation preserves the importance of local neighbors and thus makes the node representation more informative. However, due to the high sparsity of the SL graph, many genes have only a few local neighbors, and their representations aggregated from local neighbors may thus be insufficiently informative. For instance, as shown in Fig.1 (d), node *n*_4_ may learn limited feature information from its neighbors since it has only one local neighbor (i.e., *n*_6_). To tackle this issue, we consider global neighbors to further enrich the node representations. Global neighbors are defined as the nodes that have at least two hops from a given node in the graph. As shown in Fig.1 (d), nodes *n*_3_, *n*_5_, and *n*_8_ are 3-hop neighbors of node *n*_4_. Intuitively, global neighbors may contribute valuable information to the representation of the centre node. Based on this intuition, we propose a random walk with restart (RWR) (Tong *et al.*, 2008) based attention mechanism to learn node representations from global neighbors. Random walk shows powerful performance in capturing close associations between nodes and it has been successfully utilized to extract contextual information for data feature representations in multiple tasks, such as node classification (Atwood and Towsley, 2016) and recommendation systems (Zhang *et al.*, 2019). Formally, RWR is defined as follows:

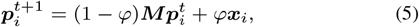

where ***M*** is the transition probability matrix obtained by normalizing the adjacency matrix *A* and *ϕ* is the restart probability, which is empirically set to 0.9. ***x***_*i*_ ∈ R^*n*^ is the initial probability vector of the *i*-th node and ***x***_*ij*_ = 1 if *j* = *i*, otherwise 0. 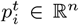 implies the probabilities of reaching other nodes at the time *t* starting from the *i*-th node, and we take 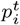 at steady state as the walking score vector for the *i*-th node. After RWR, we prioritize all SL pairs according to their walking scores and select the top *m* genes as its global neighbors for a given node. In the experiments, *m* is set to the number of local neighbors, i.e., 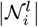.

After determining the global neighbors, we first learn the importance of the global neighbors by calculating the following coefficients 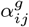:

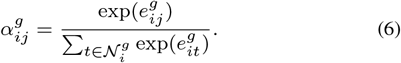

where 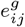 is the attention score calculated in Eq.1 and 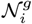 denotes the set of global neighbors of node *v*_*j*_.

Eventually, we can attain the global representation 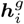 for node *v*_*i*_ by integrating the representations of global neighbors as follows:

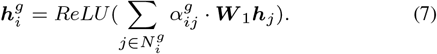

where ***W***_1_ is shared by local and global neighbors.

Similarly, we also apply the multi-head attention and repeat the node attention for *K* times. We concatenate the *K* learned representations as the final global representation as follows:

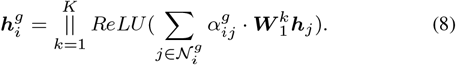

#### 2.3.3 Neural representation propagation

We have derived three different genres of representations for node *ν*_*i*_, i.e., the direct representation ***h***_i_, the local representation 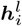 and the global representation 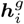. The three representations contain different semantic information. We further design a neural representation propagation module with a bi-interaction aggregator to more accurately combine these representations, which encodes the feature interactions among ***h***_*i*_ 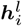 and 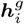 through two multilayer perceptrons (MLPs). Specifically, the bi-interaction aggregator is formulated as follows:

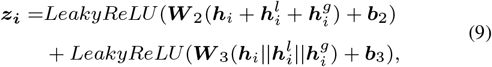

where 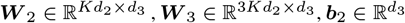, and 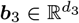 are trainable weight matrices and bias vectors respectively, and || is the concatenation operation. Here we feed ***h***_*i*_, 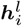 and 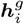 into the MLPs and consider the output ***z*_*i*_** of the last layer as the feature-specific representation of node *ν*_*i*_ as shown in Fig.1(b).

### 2.4 Feature-level attention for representation aggregation

With different feature graphs as inputs, we derive multiple feature-specific representations for each node. Since different feature graphs contain distinct context information, we further implement a feature-level attention to aggregate feature-specific r epresentations. T his p rocess f ocuses on more important feature graphs by assigning greater weight values to corresponding learned representations. We first t ransform t he feature-specific representations through a linear transformation (i.e., MLP), and then evaluate the importance of each graph by the similarity of the transformed representation and a trainable weight vector ***q*** in Eq.10. Specifically, the attention score 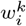 represents the importance of the *k*^*th*^ feature graph to the node *ν*_*i*_.

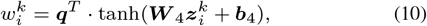

where 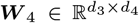 and 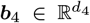 are a trainable weight matrix and a bias vector respectively, and *d*_4_ is the number of neurons in the MLPs. Note that all the matrices are shared for different feature-specific representations. To make coefficients comparable across different feature graphs, we normalize the attention scores using the softmax function:

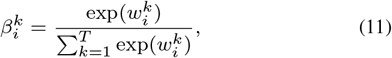

where *T* represents the number of feature graphs, and 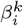 is the normalized attention score which shows the importance of the *k*^*th*^ feature graph to *ν*_*i*_.

Subsequently, we obtain the final representation *z*_*i*_ for *ν*_*i*_ by aggregating its feature-specific representations 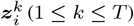 according to the normalized attention scores 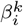 as follows:

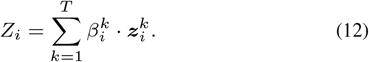

### 2.5 Optimization for SL graph reconstruction

After deriving the representation matrix *Z* for all the genes, we can then reconstruct the SL interaction network. Mathematically, we reconstruct the adjacency matrix for SL interactions in Eq. 13 and derive the reconstruction loss in Eq. 14.

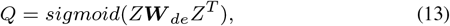

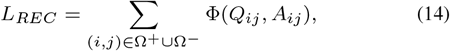

where ***W***_*de*_ is learnable latent factor that projects representations back to original feature space for genes, *sigmoid* is the activation function and *Q* is the probability score matrix with each entity representing the probability of a pair of genes to be an SL. In addition, Φ is the MSE loss (i.e., mean square error). Ω^+^ and Ω^*−*^ represent the sets of positive and negative samples for model training, respectively. To limit the influence of ***W***_*de*_ on our model, we add a constraint in Eq.14. Hence, the overall loss is defined as follows:

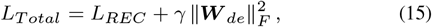

where *γ* is weight factor that is used to control the impact of ***W***_*de*_. In this work, following Long *et al.* (2020), we use the Adam optimizer (Kingma and Ba, 2015) for the optimization. Consequently, we use the reconstructed probability scores in *Q* to prioritize all the predicted SL interactions and determine the most likely SL for each gene.

## 3 Results

In this section, we first introduce the experimental setups and then demonstrate the performance of our proposed GCATSL model through both the comparison with various baselines and ablation study.

### 3.1 Experimental setups

#### 3.1.1 Datasets

The SL interactions used in our experiments were derived from database SynLethDB (Guo *et al.*, 2016) and its latest version SynLethDB-v2.0 (http://synlethdb.sist.shanghaitech.edu.cn/v2). In particular, SynLethDB was released in 2015 and it has 19,667 SL interactions involving 6,375 genes. Among 19,667 interactions, 5,740 SL interactions are computationally predicted by DAISY, 1,280 SL interactions are determined by text mining, and the rest are determined by large-scale screening techniques, such as shRNA screening and RNAi screening. SynLethDB-v2.0 has 36,741 SL interactions involving 10,218 genes, including 7,053 predicted SL interactions. We also collected a breast cancer-specific dataset from SynLethDB-v2.0, where 629 SL interactions between 612 genes are included. Table 1 summarizes the details of these three datasets. Density is the ratio of the number of existing SL interactions to the number of total possible SL interactions in the graph.

**Table 1.**
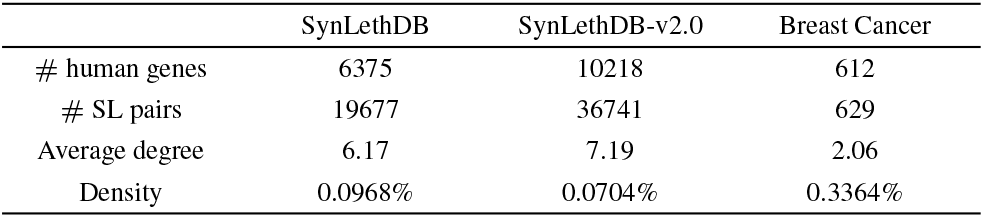
The statistics for each SL interaction dataset.

|  | SynLethDB | SynLethDB-v2.0 | Breast Cancer |
| --- | --- | --- | --- |
| # human genes | 6375 | 10218 | 612 |
| # SL pairs | 19677 | 36741 | 629 |
| Average degree | 6.17 | 7.19 | 2.06 |
| Density | 0.0968% | 0.0704% | 0.3364% |

We derived the feature graphs from Gene Ontology (GO) and PPI data. We first downloaded the ontology and annotation files from http://geneontology.org/. Two semantic similarity matrices could be calculated as feature graphs for genes based on the sub-ontologies “biological process (BP)”, and “cellular component (CC)”. We further downloaded the PPI data from BioGrid (Oughtred *et al.*, 2019) to construct a PPI network (i.e., the adjacency matrix) as the third feature graph. Note that all the SL pairs curated in this BioGrid PPI network were removed.

#### 3.1.2 Experimental settings

In this work, we conduct standard 5-fold cross-validation (CV) under the following two different settings:

- CVS1 (CV for SL pairs): random known entries in the adjacency matrix *A* (i.e., SL pairs) are sampled for testing.
- CVS2 (CV for new genes): random rows in the adjacency matrix *A* are blinded for testing.

For CVS1, we randomly divide the known SL pairs into five groups. For each round, we select in turn one group of SL pairs as positives with an equal-sized batch of randomly sampled unknown pairs as negatives for model testing. And the remaining four groups of SL pairs together with the same number of unknown pairs are used for model training. Similarly, for CVS2, we randomly select 20% rows as test samples leaving the rest of rows as training samples. Note that here negative SL pairs are randomly sampled from unknown pairs. It is common to consider all unknown pairs as negative samples for model training and testing (Cai *et al.*, 2020; Huang *et al.*, 2019; Liu *et al.*, 2019). However, the number of negative samples is too big to run traditional feature-based methods (such as SVM and Random Forest). Therefore, we follow the study (Huang *et al.*, 2020) and sample the same numbers of positive and negative samples to test the performance of various methods in our experiments. To evaluate the performance of our model, we adopt two well-known metrics that are extensively employed for link prediction, namely, area under ROC curve (AUC) and area under precision-recall curve (AUPR). To offset the bias of random splits, we repeat each experiment for 10 times and treat the average over the 10 repetitions as the final AUC and AUPR scores. Note that the blinded rows under CVS2 refer to the new genes without any known SL pairs. Therefore, CVS2 is designed to assess the prediction ability of our proposed model for new genes.

For CVS2, nodes selected for testing are blinded and thus have no local neighbors in the SL network. Hence, RWR-based sampling strategy cannot extract global neighbors for such nodes based on the SL network. To deal with this issue and make accurate predictions for new genes, here we extract local and global neighbors for testing nodes (i.e., new genes) from the PPI network while selecting local and global neighbors for training nodes from the SL network.

In our model, the training epoch was set to 600 and our model was optimized with learning rate as 0.005. The dropout rate was 0.3. In node-level attention, the number of hidden units per head was selected as 8. In RWR, we defined the restart probability *ϕ* as 0.9 and the iteration number was set to 1000. While the above parameters are empirically set, the influences of several other important parameters will be discussed in the next section, including the dimension of input raw features *d*_1_, dimension of node representation *d*_2_, number of heads *K* and weight factor *γ*.

#### 3.1.3 Baseline methods

To evaluate the performance of our model, we compare GCATSL with four state-of-the-art methods, which are recently proposed for SL prediction.

- DDGCN (Cai *et al.*, 2020) is a GCN based method developed for SL interaction prediction.
- GRSMF (Huang *et al.*, 2019) proposes a graph regularized self-representative matrix factorization algorithm for SL prediction.
- SL^2^MF (Liu *et al.*, 2019) utilizes logistic matrix factorization to learn gene representations, which are then used to identify potential SL association.
- CMF (Liany *et al.*, 2020) presents a collective matrix factorization-based method to predict SL interactions by incorporating heterogeneous data sources.

All the above methods work on automatic gene representation learning for SL prediction. Note that for all the baseline methods we adopt the default parameter values from their original implementations. Besides, we utilize the same feature graphs (i.e., two GO semantic similarity matrices from BP and CC and a PPI matrix) for all these methods for fair comparison.

We also compare our proposed GCATSL model with 10 state-of-the-art feature-based classification algorithms. They include K-Nearest Neighbors (KNN), Support Vector Machine (SVM), Random Forest (RF), Decision Tree (DT), Naive Bayesian (NB), AdaBoost, GradientBoost, Bootstrap aggregating (Bagging), MNMC (Pandey *et al.*, 2010) and MetaSL (Wu *et al.*, 2014). In particular, both MNMC and MetaSL were developed to integrate the prediction results from other eight classification approaches. We extract 18 features from various data sources (e.g., GO, PPI and etc.). Please refer to Table S1 in our supplementary materials for more details.

### 3.2 Performance evaluation

#### 3.2.1 Results on SynLethDB and SynLethDB-v2.0

We evaluate the prediction performance of our model against various baseline methods on both SynLethDB and SynLethDB-v2.0 under different CV settings as shown in Table 2.

**Table 2.** Performance comparison between baseline methods and our method on SynLethDB and SynLethDB-v2.0 under CVS1 and CVS2 settings. The best results are marked in bold and the second best is underlined.

| Methods | SynLethDB |  |  |  | SynLethDB-v2.0 |  |  |  |
| --- | --- | --- | --- | --- | --- | --- | --- | --- |
|  | CVS1 |  | CVS2 |  | CVS1 |  | CVS2 |  |
|  | AUC | AUPR | AUC | AUPR | AUC | AUPR | AUC | AUPR |
| CMF | 0.8367±0.0035 | 0.8518±0.0023 | 0.6894±0.0528 | 0.7488±0.0529 | 0.8004±0.0041 | 0.8322±0.0038 | 0.7177±0.0421 | 0.7846±0.0542 |
| SL<sup>2</sup>MF | 0.8454±0.0109 | 0.8986±0.0059 | 0.7017±0.0433 | 0.7537±0.0564 | 0.7849±0.0040 | 0.8647±0.0036 | 0.7958±0.0358 | <u>0.8212±0.0372</u> |
| GRSMF | <u>0.8853±0.0021</u> | <u>0.9187±0.0006</u> | 0.6957±0.0348 | 0.7724±0.0358 | <u>0.9116±0.0024</u> | <u>0.9361±0.0017</u> | 0.7431±0.0264 | 0.7935±0.0256 |
| DDGCN | 0.8796±0.0080 | 0.9161±0.0046 | — | — | 0.8497±0.0021 | 0.8992±0.0025 | — | — |
| RF | 0.8516±0.0018 | 0.8807±0.0020 | 0.7771±0.0366 | 0.7997±0.0447 | 0.8560±0.0051 | 0.8757±0.0044 | 0.7811±0.0194 | 0.7926±0.0248 |
| DT | 0.7206±0.0014 | 0.7907±0.0011 | 0.6645±0.0231 | 0.7278±0.0204 | 0.7181±0.0013 | 0.7888±0.0010 | 0.6604±0.0154 | 0.7255±0.0156 |
| NB | 0.7263±0.0014 | 0.7522±0.0014 | 0.7235±0.0236 | 0.7308±0.0222 | 0.7563±0.0017 | 0.7677±0.0034 | 0.7523±0.0237 | 0.7450±0.0242 |
| SVM | 0.7388±0.0007 | 0.7633±0.0025 | 0.7333±0.0226 | 0.7396±0.0205 | 0.7671±0.0048 | 0.7787±0.0045 | 0.7607±0.0185 | 0.7554±0.0139 |
| KNN | 0.7208±0.0015 | 0.7492±0.0020 | 0.6933±0.0103 | 0.7018±0.0101 | 0.7403±0.0020 | 0.7576±0.0018 | 0.7244±0.0129 | 0.7247±0.0108 |
| Bagging | 0.8512±0.0025 | 0.8803±0.0026 | 0.7714±0.0344 | 0.7982±0.0374 | 0.8556±0.0026 | 0.8761±0.0023 | 0.7757±0.0212 | 0.7867±0.0267 |
| AdaBoost | 0.7978±0.0020 | 0.8274±0.0016 | 0.7617±0.0283 | 0.7764±0.0302 | 0.7945±0.0031 | 0.8145±0.00022 | 0.7772±0.0213 | 0.7809±0.0181 |
| GradientBoost | 0.8383±0.0008 | 0.8658±0.0009 | 0.7897±0.0221 | 0.8107±0.0223 | 0.8273±0.0018 | 0.8441±0.0015 | 0.7962±0.0182 | 0.8008±0.0154 |
| MNMC | 0.8211±0.0028 | 0.8374±0.0047 | 0.7668±0.0237 | 0.7786±0.0232 | 0.8285±0.0035 | 0.8453±0.0040 | 0.7743±0.0144 | 0.7789±0.0153 |
| MetaSL | 0.8683±0.0001 | 0.8924±0.0004 | <u>0.8004±0.0243</u> | <u>0.8189±0.0243</u> | 0.8650±0.0020 | 0.8786±0.0034 | <u>0.8109±0.0155</u> | 0.8137±0.0138 |
| GCATSL | **0.9375±0.0024** | **0.9483±0.0018** | **0.8322±0.0247** | **0.8673±0.0243** | **0.9350±0.0022** | **0.9480±0.0017** | **0.8232±0.0250** | **0.8359±0.0332** |

For CVS1, we can observe that our proposed GCATSL model achieves better performance than other baseline methods. On SynLethDB, GCATSL achieves an average AUC of 0.9375 and average AUPR of 0.9483, which are 5.90% and 3.22% higher than that of the second best method GRSMF. On SynLethDB-v2.0, GCATSL achieves an average AUC of 0.9350 and average AUPR of 0.9480, which are also 2.57% and 1.27% higher than that of the runner-up GRSMF.

For CVS2, we simulate the SL prediction for new genes by randomly blinding the rows in the adjacency matrix *A* for testing. As shown in Table 2, GCATSL attains the best AUC value of 0.8322 and AUPR value of 0.8673 on SynLethDB, which are 3.97% and 5.91% better than the second best method MetaSL. We can obtain the similar conclusion from the comparison results on SynLethDB-v2.0. Note that DDGCN is not able to generate the results under CVS2, because DDGCN uses the adjacency matrix *A* of SL graph as the feature inputs and thus the testing genes have no features under CVS2 to generate predictions. In addition, GRSMF is the second best performer under CVS1, but it loses to traditional feature-based methods (e.g., GradientBoost and MetaSL) under CVS2. These results indicate that traditional feature-based methods are more robust for SL prediction with new genes. Our GCATSL combines the features extracted from other data sources with the local and global representations learned in SL graph. Therefore, GCATSL outperforms all the baseline methods for novel SL predictions under the different settings.

In our experiments, negative samples (i.e., unknown pairs) are actually unlabelled samples, and the ground truth has the information about positive labels but no information about negative labels. Therefore, we also consider the metrics that focus on evaluating the predicted positive pairs. Table S2 in our supplementary materials show the results of Recall@k (k=1000 and 5000) on both SynLethDB and SynLethDB-v2.0 under CVS1. All the above results demonstrate once again that our GCATSL performs better than state-of-the-art methods consistently.

Lastly, we further analyzed the running time of different methods. All the methods were implemented on Windows 10 operating system with a HP Z4 G4 workstation computer of an Intel W-2133 8 cores, 3.6GHz CPU, and 32G memory. Table 3 shows the running time of various methods on dataset SynLethDB, from which we can observe that our GCATSL model takes much less time than CMF, DDGCN and GRSMF, while SL2MF performs the best in terms of running time. Given that our workstation only has CPU processors, we can expect that it will take much less time to run various methods on GPU servers.

**Table 3.**
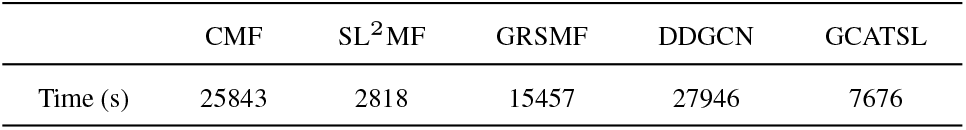
The running time of different methods on SynLethDB.

#### 3.2.2 Results on breast cancer data

To validate the effectiveness of our proposed model for specific cancer type, we implement GCATSL, four representation learning methods, and the best traditional feature-based method (i.e., MetaSL) on breast cancer data to compare their performance. Table 4 shows the comparison results of various methods under CVS1. Our method achieves higher performance than five baseline methods in terms of AUC and AUPR, demonstrating GCATSL can be successfully applied for specific cancer type.

**Table 4.** Performance comparison on the Breast Cancer. The best results are marked in bold and the second best is underlined.

| Method | AUC | AUPR |
| --- | --- | --- |
| CMF | 0.7287±0.0191 | 0.7284±0.0316 |
| SL<sup>2</sup>MF | 0.6203±0.0477 | 0.6838±0.0582 |
| GRSMF | 0.8702±0.0371 | 0.9119±0.0292 |
| DDGCN | 0.7975±0.0745 | 0.8150±0.0799 |
| MetaSL | <u>0.9103±0.0191</u> | <u>0.9151±0.0213</u> |
| GCATSL | **0.9250±0.0079** | **0.9226±0.0127** |

### 3.3 Results on DepMap data

Note that we considered randomly sampled unknown pairs as negative samples to generate all the results in Section 3.2. Some unknown pairs may be true SL pairs. Motivated by Deng *et al.* (2019), we leveraged DepMap data (Tsherniak *et al.*, 2017; McFarland *et al.*, 2020), where the effects of genes in different cell lines are recorded, to more accurately define negative SL interactions. In particular, we first downloaded CRISPR (Avana) Public 20Q4 data and then calculated co-dependency coefficients between genes based on these data using Pearson Correlation. For a given gene, we defined the top-100 genes which have the lowest co-dependency coefficients with this gene as its negative SL partners. After mapping the gene names in databases SynLethDB and DepMap, we finally obtained 275,557 negative SL interactions.

On SynLethDB dataset, we have 19,677 positive SL pairs. We further randomly sampled the same number of negative SL interactions (i.e., sampled 19,677 out of 275,557 negative SL pairs) for model training and testing. Table 5 shows the performance of various methods on SynLethDB with negative SL data defined by DepMap. It can be found that our GCATSL consistently achieves better performance than baseline methods in terms of AUC and AUPR. Compared with the results in Table 2, we observe that negative SL pairs extracted from DepMap can improve the performance of various methods including GRSMF, MetaSL and our GCATSL, demonstrating that DepMap can provide more valuable genetic co-dependency information to define high-quality negative SL data.

**Table 5.**
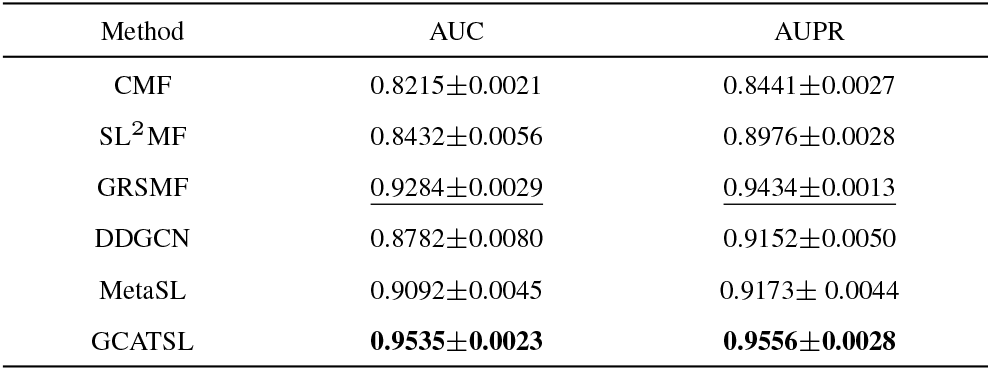
Performance comparison under CVS1 on SynLethDB with negative SL data defined by DepMap. The best results are marked in bold and the second best is underlined.

| Method | AUC | AUPR |
| --- | --- | --- |
| CMF | 0.8215±0.0021 | 0.8441±0.0027 |
| SL<sup>2</sup>MF | 0.8432±0.0056 | 0.8976±0.0028 |
| GRSMF | <u>0.9284±0.0029</u> | <u>0.9434±0.0013</u> |
| DDGCN | 0.8782±0.0080 | 0.9152±0.0050 |
| MetaSL | 0.9092±0.0045 | 0.9173±0.0044 |
| GCATSL | **0.9535±0.0023** | **0.9556±0.0028** |

### 3.4 Ablation study

Our GCATSL consists of a two-level attention mechanism, i.e., node-level attention and feature-level attention. To evaluate their impact on our model, we derive two model variants GCATSL-N and GCATSL-F, which refer to our model using node-level attention only and feature-level attention only, respectively. For example, GCATSL-N uses average weights instead of the feature-level attention scores in Eq.12. Fig.2 (a) shows that GCATSL-N and GCATSL-F achieve worse performance than GCATSL, indicating both node-level and feature-level attention are effective in capturing different semantic information for genes. Moreover, feature-level attention looks more important than node-level attention, as GCATSL-F performs better than GCATSL-N in terms of both AUC and AUPR.

**Fig. 2.**
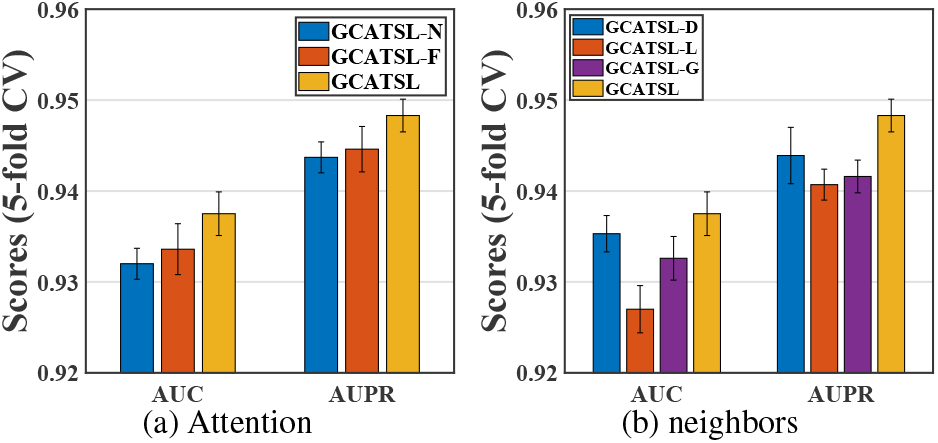
Comparison between GCATSL and its variants on SynLethDB under CVS1.

Recall that we combine three types of representations (*h*, *h*^*l*^ and *h*^*g*^) from different contexts in Eq.9. We derive the following model variants to study the importance of different contexts.

- GCATSL-D: it dose not include the direct features *h* in Eq.9.
- GCATSL-L: it dose not consider *h*^*l*^ from local neighbors.
- GCATSL-G: it dose not consider *h*^*g*^ from global neighbors.

As shown in Fig.2 (b), GCATSL achieves higher AUC and AUPR values than GCATSL-D, GCATSL-L and GCATSL-G. We can thus conclude that all three genres of context information can help improve the prediction performance of our model. Also, we can observe that local neighbors are the most important context for gene representation learning and SL prediction, as GCATSL-L which uses only *h* and *h*^*g*^ achieves the lowest performance.

Lastly, we exploit three different types of biological data sources (i.e., BP, CC and PPI) to generate feature graphs for genes as inputs of our GCATSL. Here, we examine the contribution of each data source (i.e., each feature graph). Fig.3 shows that integrating all the three data sources help our model to achieve the highest prediction performance. Moreover, GO similarity (i.e., BP and CC) contributes more than PPI while BP can bring relatively more informative features than CC.

**Fig. 3.**
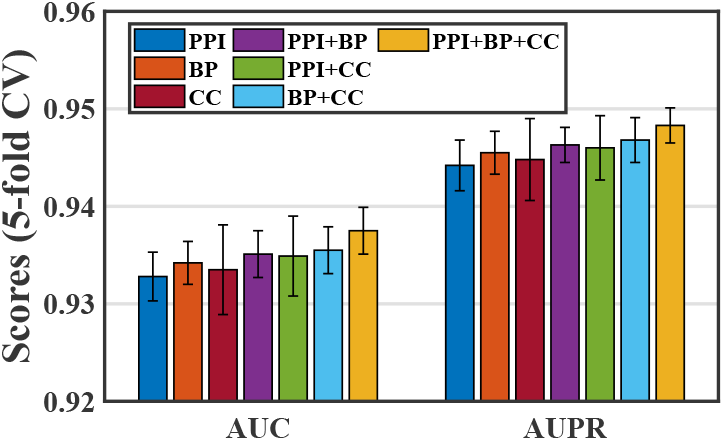
Effect of different feature graphs on GCATSL.

### 3.5 Parameter analysis

There are several important parameters that influence the performance of our model, such as dimension of input raw features *d*_1_, dimension of node representation *d*_2_, number of heads *K* and weight factor *γ*. In this section, we perform the sensitivity analysis for these parameters. Note that here all the experiments are conducted on SynLethDB under the CVS1 setting.

We implemented PCA on feature graphs to filter out the noises and reduced the feature dimension. Thus the output size of PCA is the input feature dimension *d*_1_ of our GCATSL model. We then evaluate our model by picking *d*_1_ from {8, 16, 32, 64, 128, 256}. Fig.4 (a) shows that as *d*_1_ varies, the performance first gradually increases and then decreases, with *d*_1_ = 128 achieving its best performance. In our model, the number of neurons of MLP in Fig.1 determines the dimension of node representation *d*_2_. To measure the impact of *d*_2_ on our model, we choose its value from {4, 8, 16, 32, 64, 128}. As shown in Fig.4 (b), we can observe that our model is slightly influenced by *d*_2_, and attain the best performance when *d*_2_ is equal to 64. In the node-level attention, we adopt multi-head attention mechanism to obtain more informative node representation. By varying the number of heads *K* from 1 to 8 with a step value of 1, we can observe that our model is relatively robust as both AUC and AUPR are quite stable when *K* varies in Fig.4 (c). In addition, we use weight factor *γ* in Eq.15 to regularize the influence of weight matrix ***W***_*de*_. To evaluate its impact, we choose its value from {0.0001, 0.0005, 0.001, 0.005, 0.01, 0.05}. As shown in Fig.4 (d), our model is stable when *γ* is set to small values. However, GCATSL achieves low performance when *γ* is larger than 0.001 and the best performance is obtained when *γ* is set to 0.0005.

**Fig. 4.**
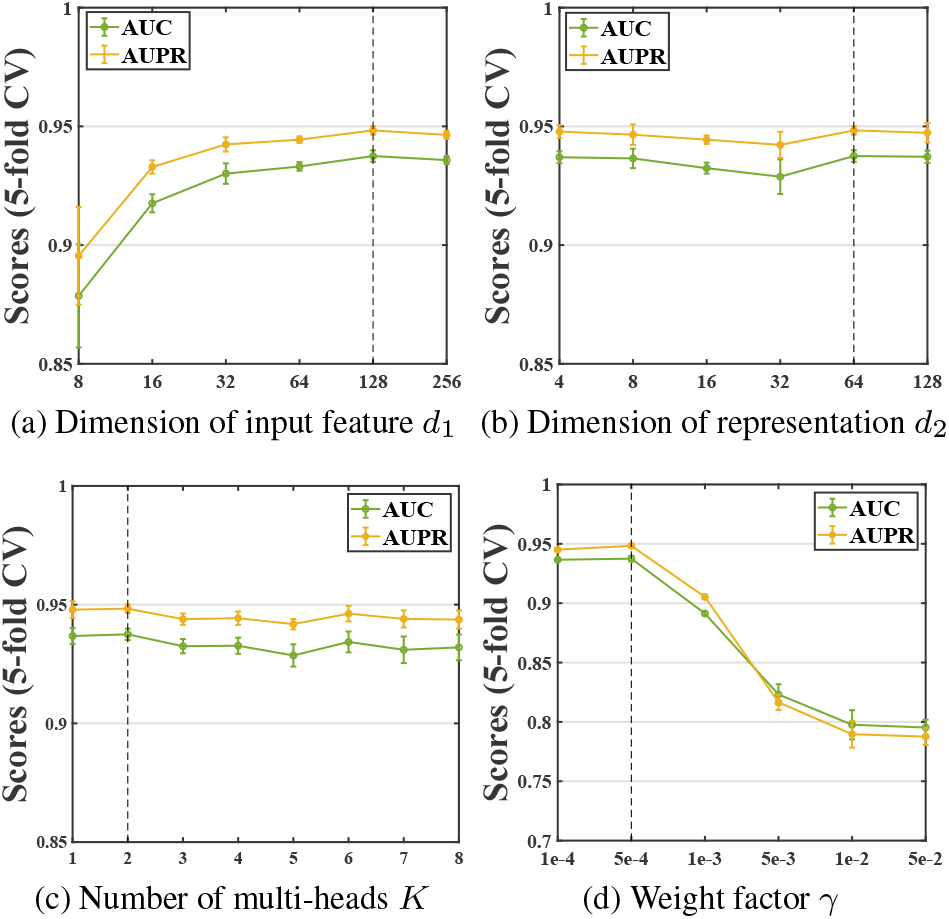
Parameter sensitivity analysis for GCATSL on SynLethDB under CVS1.

### 3.6 Case study

To further validate the performance of the GCATSL model, we conduct a case study based on dataset SynLethDB. More specifically, we utilize all the known SL pairs in SynLethDB as training samples to train the model, then prioritize all the unknown SL pairs according to their predicted scores. We check how many unknown SL pairs among the top 1000 pairs are reported in SynLethDB-v2.0 and supported by biomedical literature.

Our results show that 36 out of top 1000 unknown pairs are confirmed in SynLethDB-v2.0. Table 6 shows 20 selected SL pairs, while the rest of SL pairs are displayed in supplementary materials. The validation evidence in Table 6 is available in SynLethDB-v2.0. We find that many pairs are validated by wet-labs. For example, the pairs between KRAS and DDR1, BID and SSH3, are confirmed by shRNA screening (Vizeacoumar *et al.*, 2013). Row 4 (MYC and NTRK1), row 7 (CYP1B1 and KRAS) and row 8 (E2F1 and KRAS) are verified by siRNA screening technology (Luo *et al.*, 2009). SL pairs between KRAS and PIK3CA/TBL1XR1/SRP9/LUC7L2 are detected by CRISPR screening (Martin *et al.*, 2017). In addition, there are 4 SL pairs that are supported by other in silico methods. For example, row 1 (BCR and KRAS), row 3 (KRAS and RET) and row 17 (KRAS and MSH2) were predicted by Chang *et al.* (2016). In fact, KRAS is the most commonly mutated oncogene in human cancer, and is considered as a high-priority therapeutic target. Table 6 shows that we predict several SL partners for KRAS.

**Table 6.** 20 SL pairs confirmed in SynLethDB-v2.0 among the top 1000 predicted pairs.

| No. | Gene 1 | Gene 2 | PubMed ID | Evidence | No. | Gene 1 | Gene 2 | PubMed ID | Evidence |
| --- | --- | --- | --- | --- | --- | --- | --- | --- | --- |
| 1 | BCR | KRAS | 27655641 | in-silico prediction | 11 | BID | KRAS | 24104479 | shRNA screening |
| 2 | DDR1 | KRAS | 24104479 | shRNA screening | 12 | KRAS | NHP2 | 28700943 | CRISPR screening |
| 3 | KRAS | RET | 27655641 | in-silico prediction | 13 | KRAS | SSH3 | 24104479 | shRNA screening |
| 4 | MYC | NTRK1 | 22623531 | siRNA screening | 14 | ABL1 | KIT | 26637171 | siRNA screening |
| 5 | KRAS | PIK3CA | 26627737 | CRISPR-Cas9 | 15 | NTRK1 | PDGFRB | 26637171 | siRNA screening |
| 6 | KRAS | TBL1XR1 | 28700943 | CRISPR screening | 16 | KIT | PDGFRA | 31300006 | in-silico prediction |
| 7 | CYP1B1 | KRAS | 22613949 | siRNA screening | 17 | KRAS | MSH2 | 27655641 | in-silico prediction |
| 8 | E2F1 | KRAS | 22613949 | siRNA screening | 18 | KRAS | SRP9 | 28700943 | CRISPR screening |
| 9 | ABL1 | PDGFRB | 26637171 | siRNA screening | 19 | KRAS | LUC7L2 | 28700943 | CRISPR screening |
| 10 | KIT | PDGFRB | 26637171 | siRNA screening | 20 | CDK1 | KRAS | 26881434 | siRNA screening |

Besides, we compare the performance of different methods in identifying novel SL pairs. The Fig. S1 in supplementary materials shows our model performs better than baseline methods. Therefore, it could be concluded that our proposed model is an effective and promising tool to assist pharmacologists and biologists in screening potential anticancer targets for drug developments in the future.

## 4 Discussion and Conclusion

Synthetic lethal (SL) is a promising type of genetic interaction that plays an critical role in targeted anticancer therapeutics. Considering the limitations of experimental methods, in silico methods provide a useful guide for the wet-lab experiments to screen candidate SL pairs.

In this work, we propose a novel Graph Contextualized Attention Network framework, named GCATSL, for SL prediction. First, we leveraged different biological data sources to construct multiple feature graphs for genes. We then implemented PCA on each graph to extract valuable features and reduce dimensionality. Second, we designed a hierarchical attention mechanism, i.e., node-level attention and feature-level attention, to learn node representations from multiple feature graphs. In particular, we designed node-level attention mechanism to effectively preserve the importance of local and global neighbors to learn better local and global representations for the nodes, respectively. We further introduced a neural network architecture to aggregate the original features with the local and global representations, and thus derived the feature-specific representations. Finally, we implemented feature-level attention to integrate feature-specific representations by taking the importance of different feature graphs into account. Extensive experimental results on three datasets demonstrated that our proposed GCATSL model performed better than 14 existing methods in predicting novel SL interactions for both existing genes and new genes.

While our model achieves good prediction performance, there are still some limitations expected to be overcome. First, node features in our model are manually extracted from various data sources. We would like to investigate pretraining strategies for automatic feature extraction, which has achieved great success in natural language processing applications (Devlin *et al.*, 2019). Second, in our current case study, we train the model based on the original SynLethDB database and then simply validate the predicted SL pairs based on its updated version SynLethDB-v2.0. In the future, we plan to collaborate with biologists and conduct wet-lab experiments to verify the predicted results.

## Funding

This work was supported by the National Natural Science Foundation of China [61873089], the Key Program of National Natural Science Foundation of China [62032007] and the Chinese Scholarship Council (CSC) [201906130027].

## Acknowledgements

The authors would like to gratefully acknowledge Jie Wang from ShanghaiTech University for her help on analyzing DepMap dataset.

## Notes

### Competing Interest Statement

The authors have declared no competing interest.

## References

Apaolaza, I. et al. (2017). An in-silico approach to predict and exploit synthetic lethality in cancer metabolism. Nature communications, 8(1), 1–9.

Atwood, J. and Towsley, D. (2016). Diffusion-convolutional neural networks. In Advances in neural information processing systems, pages 1993–2001.

Benstead-Hume, G. et al. (2019). Predicting synthetic lethal interactions using conserved patterns in protein interaction networks. PLoS computational biology, 15(4), e1006888.

Cai, R. et al. (2020). Dual-dropout graph convolutional network for predicting synthetic lethality in human cancers. Bioinformatics, 36(16), 4458–4465.

Chan, D. A. et al. (2011). Targeting glut1 and the warburg effect in renal cell carcinoma by chemical synthetic lethality. Science translational medicine, 3(94), 94ra70.

Chang, J.-G. et al. (2016). Uncovering synthetic lethal interactions for therapeutic targets and predictive markers in lung adenocarcinoma. Oncotarget, 7(45), 73664.

Das, S. et al. (2019). Discoversl: an r package for multi-omic data driven prediction of synthetic lethality in cancers. Bioinformatics, 35(4), 701–702.

Deng, X. et al. (2019). Sl-biodp: Multi-cancer interactive tool for prediction of synthetic lethality and response to cancer treatment. Cancers, 11(11), 1682.

Devlin, J. et al. (2019). Bert: Pre-training of deep bidirectional transformers for language understanding. In NAACL-HLT.

Du, D. et al. (2017). Genetic interaction mapping in mammalian cells using crispr interference. Nature methods, 14(6), 577.

Ezzat, A. et al. (2019). Computational prediction of drug–target interactions using chemogenomic approaches: an empirical survey. Briefings in bioinformatics, 20(4), 1337–1357.

Guo, J. et al. (2016). Synlethdb: synthetic lethality database toward discovery of selective and sensitive anticancer drug targets. Nucleic acids research, 44(D1), D1011–D1017.

Hartwell, L. H. et al. (1997). Integrating genetic approaches into the discovery of anticancer drugs. Science, 278(5340), 1064–1068.

Hong, Z. et al. (2020). Identifying enhancer–promoter interactions with neural network based on pre-trained dna vectors and attention mechanism. Bioinformatics, 36(4), 1037–1043.

Huang, J. et al. (2019). Predicting synthetic lethal interactions in human cancers using graph regularized self-representative matrix factorization. BMC bioinformatics, 20(19), 1–8.

Huang, Y.-a. et al. (2020). Graph convolution for predicting associations between mirna and drug resistance. Bioinformatics, 36(3), 851–858.

Iglehart, J. D. and Silver, D. P. (2009). Synthetic lethality–a new direction in cancer-drug development. New England Journal of Medicine, 361(2), 189.

Jacunski, A. et al. (2015). Connectivity homology enables inter-species network models of synthetic lethality. PLoS Comput Biol, 11(10), e1004506.

Jerby-Arnon, L. et al. (2014). Predicting cancer-specific vulnerability via data-driven detection of synthetic lethality. Cell, 158(5), 1199–1209.

Kingma, D. P. and Ba, J. (2015). Adam: A method for stochastic optimization. In 3rd international conference for learning representations, San Diego.

Liany, H. et al. (2020). Predicting synthetic lethal interactions using heterogeneous data sources. Bioinformatics, 36(7), 2209–2216.

Liu, Y. et al. (2016). Neighborhood regularized logistic matrix factorization for drug-target interaction prediction. PLoS computational biology, 12(2), e1004760.

Liu, Y. et al. (2019). Sl2mf: Predicting synthetic lethality in human cancers via logistic matrix factorization. IEEE/ACM Transactions on Computational Biology and Bioinformatics, 17(3), 748–757.

Long, Y. et al. (2020). Predicting human microbe-drug associations via graph convolutional network with conditional random field. Bioinformatics, 36(19), 4918–4927.

Luo, J. et al. (2009). A genome-wide rnai screen identifies multiple synthetic lethal interactions with the ras oncogene. Cell, 137(5), 835–848.

Martin, T. D. et al. (2017). A role for mitochondrial translation in promotion of viability in k-ras mutant cells. Cell reports, 20(2), 427–438.

McFarland, J. M. et al. (2020). Multiplexed single-cell transcriptional response profiling to define cancer vulnerabilities and therapeutic mechanism of action. Nature communications, 11(1), 1–15.

Natarajan, N. and Dhillon, I. S. (2014). Inductive matrix completion for predicting gene–disease associations. Bioinformatics, 30(12), i60–i68.

O’Neil, N. J. et al. (2017). Synthetic lethality and cancer. Nature Reviews Genetics, 18(10), 613–623.

Oughtred, R. et al. (2019). The biogrid interaction database: 2019 update. Nucleic acids research, 47(D1), D529–D541.

Pandey, G. et al. (2010). An integrative multi-network and multi-classifier approach to predict genetic interactions. PLoS computational biology, 6(9), e1000928.

Sinha, S. et al. (2017). Systematic discovery of mutation-specific synthetic lethals by mining pan-cancer human primary tumor data. Nature communications, 8(1), 1–13.

Srihari, S. et al. (2015). Inferring synthetic lethal interactions from mutual exclusivity of genetic events in cancer. Biology direct, 10(1), 1–18.

Tong, H., Faloutsos, C., and Pan, J.-Y. (2008). Random walk with restart: fast solutions and applications. Knowledge and Information Systems, 14(3), 327–346.

Tsherniak, A. et al. (2017). Defining a cancer dependency map. Cell, 170(3), 564–576.

Veličković, P. et al. (2018). Graph attention networks. In International Conference on Learning Representations.

Vizeacoumar, F. J. et al. (2013). A negative genetic interaction map in isogenic cancer cell lines reveals cancer cell vulnerabilities. Molecular systems biology, 9(1), 696.

Wu, M. et al. (2014). In silico prediction of synthetic lethality by meta-analysis of genetic interactions, functions, and pathways in yeast and human cancer. Cancer informatics, 13, CIN–S14026.

Zhang, C. et al. (2019). Heterogeneous graph neural network. In Proceedings of the 25th ACM SIGKDD International Conference on Knowledge Discovery & Data Mining, pages 793–803.

Zhang, F. et al. (2015). Predicting essential genes and synthetic lethality via influence propagation in signaling pathways of cancer cell fates. Journal of bioinformatics and computational biology, 13(03), 1541002.

Zhang, Z.-C. et al. (2020). A graph regularized generalized matrix factorization model for predicting links in biomedical bipartite networks. Bioinformatics, 36(11), 3474–3481.

